# Kidney single-cell transcriptomes uncover SGLT2i-induced metabolic reprogramming via restoring glycolysis and fatty acid oxidation

**DOI:** 10.1101/2023.10.31.564836

**Authors:** Ying Shi, Vivek Bhalla

## Abstract

Approximately 40% of individuals with chronic kidney disease have type 2 diabetes mellitus, and diabetic kidney disease is the leading cause of end-stage kidney disease worldwide. Inhibitors of sodium-glucose cotransporter 2 (SGLT2) have been demonstrated to be effective in glucose control, improving cardiovascular outcomes and the progression of kidney disease. However, the protective role of SGLT2 inhibition on kidney metabolism is not fully understood. To explore these mechanisms further, we conducted analysis of publicly available single-cell RNA sequencing data of db/db mice treated with an SGLT2 inhibitor(dapagliflozin) and accompanying controls. We found that proximal tubule cells exhibited impaired glycolysis and high fatty acid oxidation in diabetes compared with control mice. SGLT2 inhibition reversed this metabolic dysfunction by reducing glycolysis and its substrate accumulation. SGLT2 inhibition also upregulates high fatty oxidation without increasing the uptake of fatty acids and elongation, along with low lipotoxicity. Surprisingly, both SGLT2(+) and SGLT2(-) cells show gene consistent changes in expression of metabolic genes, consistent with a non-cell autonomous effect of dapagliflozin treatment. This study demonstrates the protective role of SGLT2 inhibition via restoring metabolic dysfunction.

## Introduction

11.3 % of the US population has been diagnosed with diabetes (1), and 96 million Americans, 38% of the adult US population, show signs of pre-diabetes (1). In the United States, 1 in 3 adults will develop diabetes sometime in their lifetime (2). Multiple glucose-lowering drugs are available to manage blood glucose for the treatment of diabetes, including insulin, metformin, sulfonylureas, glucagon-like peptide 1 (GLP-1) agonists, and sodium–glucose cotransporter 2 (SGLT2) inhibitors (3–6). Compared to other therapies, SGLT2 inhibitors (SGLT2i) demonstrate remarkable protection in preventing kidney damage progression and improving cardiovascular outcomes in large-scale randomized clinical trials (7–10).

It is well-recognized that SGLT2 is mainly expressed in the proximal tubule (PT) cells and accounts for glucose and sodium reabsorption (11, 12). Some studies reported that SGLT2 reduces intraglomerular pressure and reverses tubuloglomerular feedback via increased distal delivery of sodium (13–15). Previous studies also suggest that SGLT2 inhibition protects the kidney from renal inflammation and fibrosis (16–19). However, the renoprotective mechanisms of SGLT2 inhibition are not well elucidated.

Impaired glycolysis induces the accumulation of glucose substrates that contribute to mitochondrial dysfunction, apoptosis, and kidney pathology in diabetes (20, 21). SGLT2 activation results in increased intracellular glucose flux, providing more substrate for glycolysis (22). By contrast, SGLT2 inhibition suppresses the transcription of genes that regulate glycolysis, gluconeogenesis, and tricarboxylic acid cycle (TCA) pathways in individuals with type 2 diabetes (23). Similarly, in a type 1 diabetic mouse model, SGLT2 inhibition restores the diabetes-induced increase in expression of glycolytic enzymes hexokinase 2 and pyruvate kinase M2 (24). However, in another mouse model of type 1 diabetes, treatment with an SGLT2i failed to rescue glycolysis. SGLT2 inhibition increases glucose oxidation in multiple organs (25). Thus, the role of SGLT2i in glycolysis is still under debate. Moreover, whether the effect of these drugs is different in SGLT2(+) vs. SGLT2(-) cells has not been explored.

In the present study, we aimed to characterize the effects of SGLT2i treatment on SGLT2(+) vs. SGLT2(-) cells in a kidney mouse model of type 2 diabetes. By analyzing the scRNA-seq data set from diabetic mice treated with dapagliflozin (26), we were able to dissect the specific metabolic responses of PT cells to SGLT2i.

## Materials and Methods

### Data Acquisition

We obtained the scRNA-seq data set from the study conducted by the He laboratory (https://www.ncbi.nlm.nih.gov/geo/, accession number: GSE181382) (26). We selected 3 groups from the dataset, including two control samples (*db/m* mice), three diabetic samples (*db/db* mice at 10 weeks of age were treated with phosphate-buffered saline (PBS)), and 4 treated samples (*db/db* mice at 10 weeks of age were treated with dapagliflozin). PBS and dapagliflozin were administered daily by oral gavage for 8 weeks. All mice were sacrificed at the age of 18 weeks.

### scRNA-seq data quality control and analysis

We removed cells with genes expressed in fewer than three cells and cells expressing fewer than 200 genes, or more than 5000 genes. We also removed cells with > 50% mitochondrial genes. We used the top 2,500 highly variable genes by the “FindVariableFeatures” function with the default method.

### Unsupervised clustering and cell annotation

We selected the first 30 principal components (PCs) identified from the principal component analysis to construct the K-nearest neighbor graph using the “FindNeighbor” function, and unsupervised clustering analysis by the “FindClusters” function with the default method, with a resolution of 0.5. We used UMAP for subsequent dimensionality reduction usage using the “RunUMAP” function. We identified the cell markers of cell types by “FindAllMarkers” and used UMAP to visualize the annotation. We used the tubular cell markers (Kap, Lrp2, Slc5a2, slc7a13, slc13a3, slc13a1, slc5a12) for identifying PT cells, Umod for thick ascending limb, Ltd for T cells, S100a8 and S100a9 for neutrophils and Ly6c1 and Cd74 for macrophages.

### Pseudo-time analysis of PT cells

We selected the PT cells as a subset to further explore the mechanism using the “subset” function. We used the Monocle3 package to perform pseudo-time analysis using default parameters.

### Annotation of SGLT2+ cells and negative cells

We selected the cells with reads of the Slc5a2 gene greater than zero and annotated these cells as SGLT2+ Cells (SGLT2(+) cells). Consequently, all other PT cells were labeled as Negative Cells (SGLT2(-) cells).

### Differentially expressed genes and GO enrichment analysis

We identified the differentially expressed genes (DEGs) between the SGLT2(+) cells and SGLT2(-) cells with “FindMarkers” and genes with p ≤ 0.05 and log fold change ≥ 0.25. We matched the DEGs for the GO enrichment by ClusterProfiler 3.0.4 and GO terms with p ≤ 0.05 as statistically significant.

### Signature scores of gene sets

We generated the gene sets for glycolysis and fatty acid oxidation. The geneset of positive regulation of fatty acid metabolic processes was obtained from Mouse Genome Informatics. The combined signature score was calculated using the UCell (27). The full gene list is listed in **Table S1**.

### Gene expression visualization

We used the “VlnPlot” for visualizing SGLT2 expression between groups and the “DotPlot” for the pathways.

## Results

### Single-cell profile of control and diabetic mouse kidneys with or without SGLT2i treatment

In total, 47998 single-cell transcriptomes were generated after quality control and filtering (**Fig. 1A**). Using the UMAP algorithm for dimensionality reduction and unsupervised clustering, we identified the five major cell populations: PT cells, thick ascending limb cells, T cells, neutrophils, and macrophages (**Fig. 1B**). A full list of marker genes is provided in Methods. To explore renoprotection mechanisms of SGLT2i, we generated a subset of PT cells with 41165 single-cell transcriptomes by selecting the PT cells (**Fig. 1C** and **1D**). Using unsupervised clustering, we identified 12 clusters in the PT subset. By comparing the SGLT2 gene expression (Slc5a2) in groups, we found the diabetic group had the highest expression compared to the control group and diabetic group (P<0.001, **Fig. 1F**). In the treatment group, this increase was abrogated (P<0.0001, **Fig. 1F**). For SGLT2(+) and SGLT2(-) cells (**Fig. 1G**), the majority of cells in all treatment groups were SGLT2(-) (**Fig. 1H**). We observed the control group had a similar proportion of SGLT2(+) cells to the treatment group. In contrast, the diabetic mice had the least SGLT2(+) cells. Taken together, SGLT2i treatment suppresses the diabetes-induced elevation in SGLT2 expression in PT cells.

**Figure 1:**
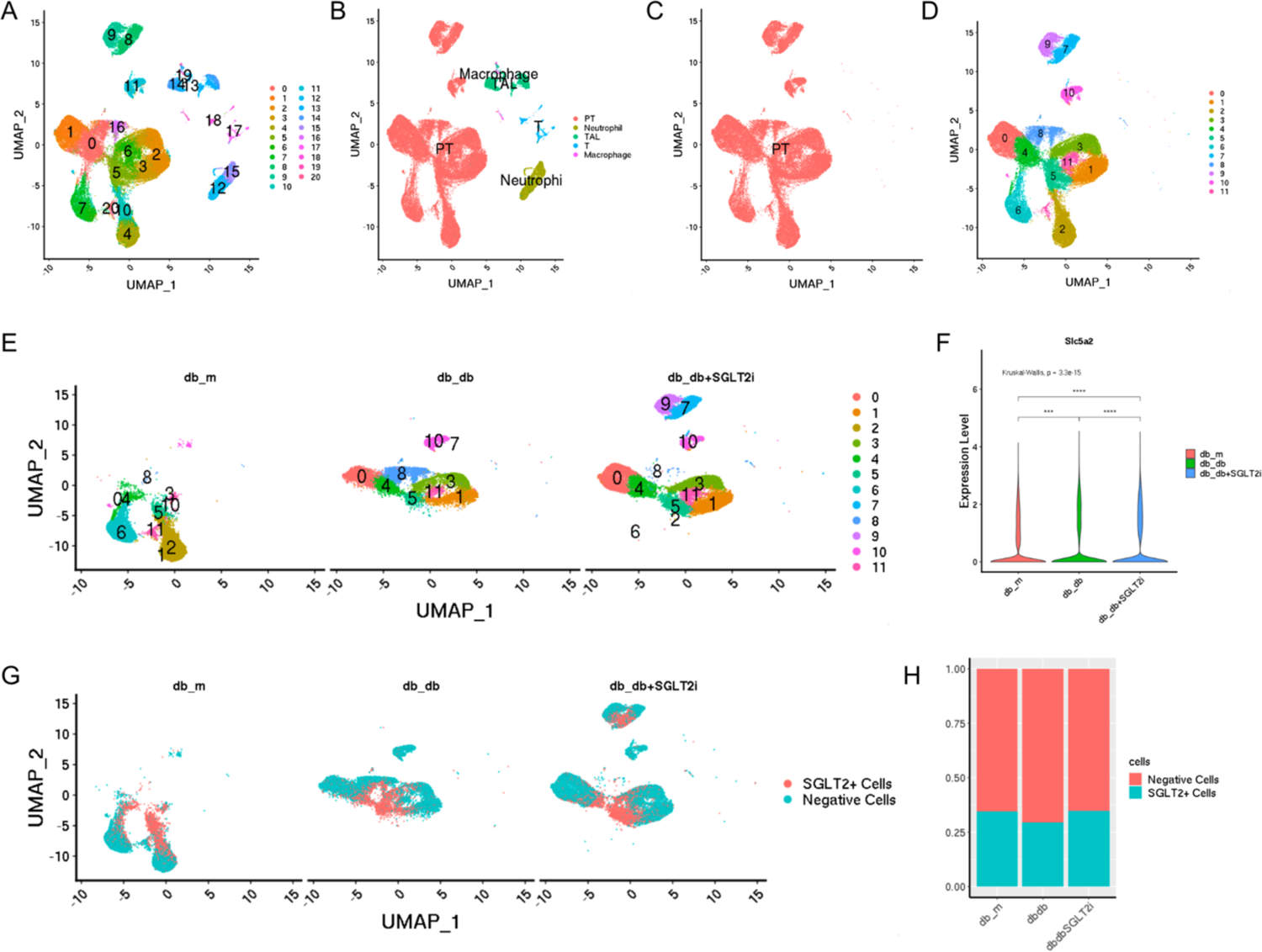
Single-cell profile of health and diabetic mouse kidneys with or without sodium-glucose cotransporter 2 (SGLT2) inhibitor treatment. **(A),** UMAP representation of all cells of scRNA-seq. **(B),** Cell annotation. **(C),** UMAP representation of proximal tubular (PT). **(D),** Clusters of PT cells. **(E),** Cluster distribution in groups. **(F),** SGLT2 expression in groups. **(G),** Distribution of SGLT2(+) (SGLT2+ Cells) and SGLT2(-) cells (Negative Cells) in groups. **(H),** Fractions of SGLT2(+) and SGLT2(-) cells of each group. All differences were analyzed by the Wilcox test. ***, P < 0.001. ****, P < 0.0001.

### SGLT2i treatment reduced glycolysis in diabetes

To assess the activation of SGLT2 in diabetes and the inhibition of SGLT2i treatment, we analyzed the scores of glycolysis in all groups and SGLT2(+) vs. SGLT2(-) cells using signature scoring analysis. Glycolysis was significantly higher in the diabetic group compared to the control group and was decreased in the treatment group (p<0.0001, **Fig 2A**). As shown in Fig. 2B, we observed a similar increase of glycolysis in the diabetic group in both SGLT2(+) vs. SGLT2(-) cells, while the SGLT2i treatment reversed the elevation (P<0.0001, **Fig 2B**).

**Figure 2:**
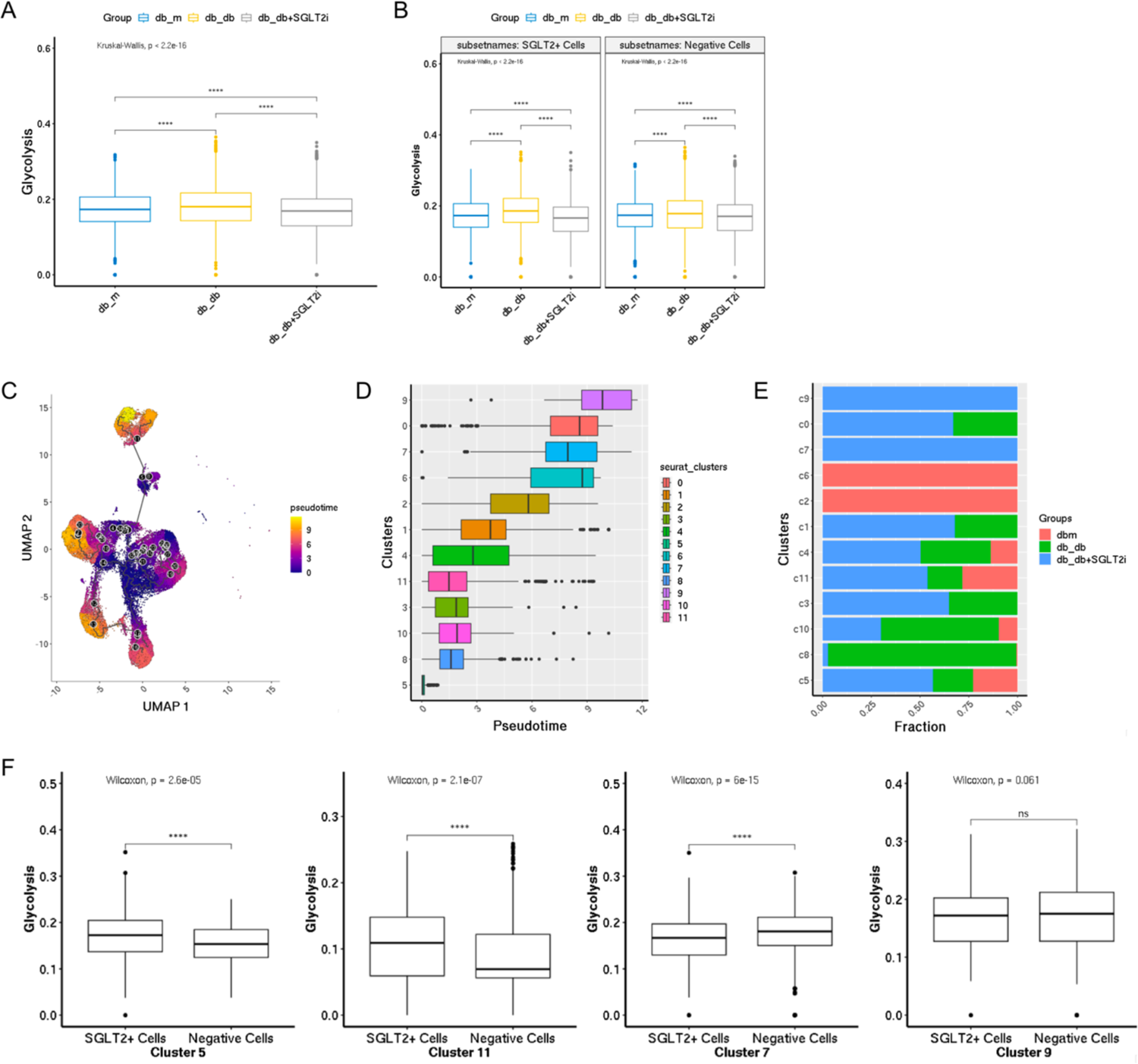
Glycolysis in diabetes. **(A),** Signature scoring analysis of glycolysis in groups. **(B),** Signature scoring analysis of glycolysis in SGLT2(+) vs. SGLT2(-) cells in groups. **(C),** Trajectory of PT cells. **(D),** Differentiation trajectory by ordered clusters. **(E),** Fractions of each group of each cluster. **(F),** Signature scoring analysis of glycolysis in clusters 5, 11, 7, 9. All differences were analyzed by the Wilcox test. ns, P>=0.05. ****, P < 0.0001.

As shown in **Fig. 1E**, the distribution of PT cells of groups had different patterns in the UMAP. Most cells in the control group (*db/m*) belonged to clusters 2 and 6, while the diabetic group (*db/db*) had more cells in clusters 8 and 10, and the treatment group (*db/db*+SGLT2i) had more cells in clusters 7 and 9. To investigate cell differentiation under diabetes and treatment with SGLT2i, we next conducted the cell trajectory analysis (**Fig. 2C**). As shown in **Fig. 2E**, cluster 5 was the starting point of cell differentiation and cluster 9 was the end. The early differentiated clusters 8 and 10 had more cells from the diabetic group, while the control group had more cells in the later differentiated clusters 2 and 6, and the treatment group occupied the majority in clusters 7, 0, and 9 (**Fig. 2E**). In addition, most early differentiated clusters contained cells from all groups (**Fig. 2D** and **2E**). In contrast, cells in clusters 2, 6, 7, and 9 came from either the control group or the treatment group (**Fig. 2E**). Next, we scored the glycolysis in clusters 5, 11, 7, and 9 between SGLT2(+) vs. SGLT2(-) cells according to the direction of cell differentiation (**Fig. 2F**). In cluster 5, SGLT2(-) cells showed lower glycolysis than SGLT2(+) cells, similar to cluster 11. However, the difference between SGLT2(+) vs. SGLT2(-) cells was reduced, and no difference was found in cluster 9. These data demonstrate that SGLT2i treatment decreased glycolysis in diabetes in both SGLT2(+) and SGLT2(-) cells. This decrease in SGLT2(-) cells was more significant in the early stages of differentiation.

Next, we further analyzed how SGLT2i treatment regulated glycolysis. We compared the expression levels of glycolytic enzymes (**Fig. 4A** and **4B**). Compared to the control group, the diabetic group had higher expression of Hk1, Pfkl, Pgk1, and Pkm, which induce irreversible reactions in glycolysis(28) (**Fig. 4A**). The diabetic group also showed lower expression levels of Mpc1/2 and Pdk3/4, indicating less pyruvate transporting to mitochondria for fueling the TCA cycle. In contrast, the treatment group showed reversed expression levels of Hk1, Pfkl, Pgk1, and Pkm, and higher expression of Mpc1/2 and Pdk4 compared to the diabetic group (**Fig. 4B**).

### SGLT2i treatment elevated fatty acid oxidation in diabetes without increasing lipotoxicity

To study the mechanism, we computed the DEGs of cluster 9 and cluster 7 between the SGLT2(+) vs. SGLT2(-) cells and mapped these DEGs to the database of biological processes (**Fig. 3A**). We found the top biological process in both clusters was the fatty acid metabolic process (**Fig.3B**). To explore the mechanism, we scored the gene set of fatty acid oxidation. We found that the diabetic group had increased fatty acid oxidation compared to the control group and the treatment further increased the oxidation (**Fig. 3C**). As shown in **Fig. 3C** and **3D**, a similar trend was found in both SGLT2(+) vs. SGLT2(-) cells. We next scored the fatty acid oxidation based on cell differentiation. These data showed no difference or little difference between SGLT2(+) vs. SGLT2(-) cells in early differentiated clusters 5 (**Fig. 3E**). However, the SGLT2(-) cells had significantly higher fatty acid oxidation compared to SGLT2(+) cells in clusters 11 but lower oxidation in cluster 7 and 9 (**Fig. 3E**), suggesting that SGLT2i treatment increased fatty acid oxidation differently in different differentiated stages. By the end of the treatment, SGLT2(+) cells exhibited higher fatty acid oxidation than SGLT2(-) cells (p<0.0001, **Fig. 3E**). Consistently, compared to the diabetic group, SGLT2i treatment further increased the positive regulation of the fatty acid oxidation, in both SGLT2(+) vs. SGLT2(-) cells (P<0.0001, **Fig. 3F** and **3G**).

**Figure 3:**
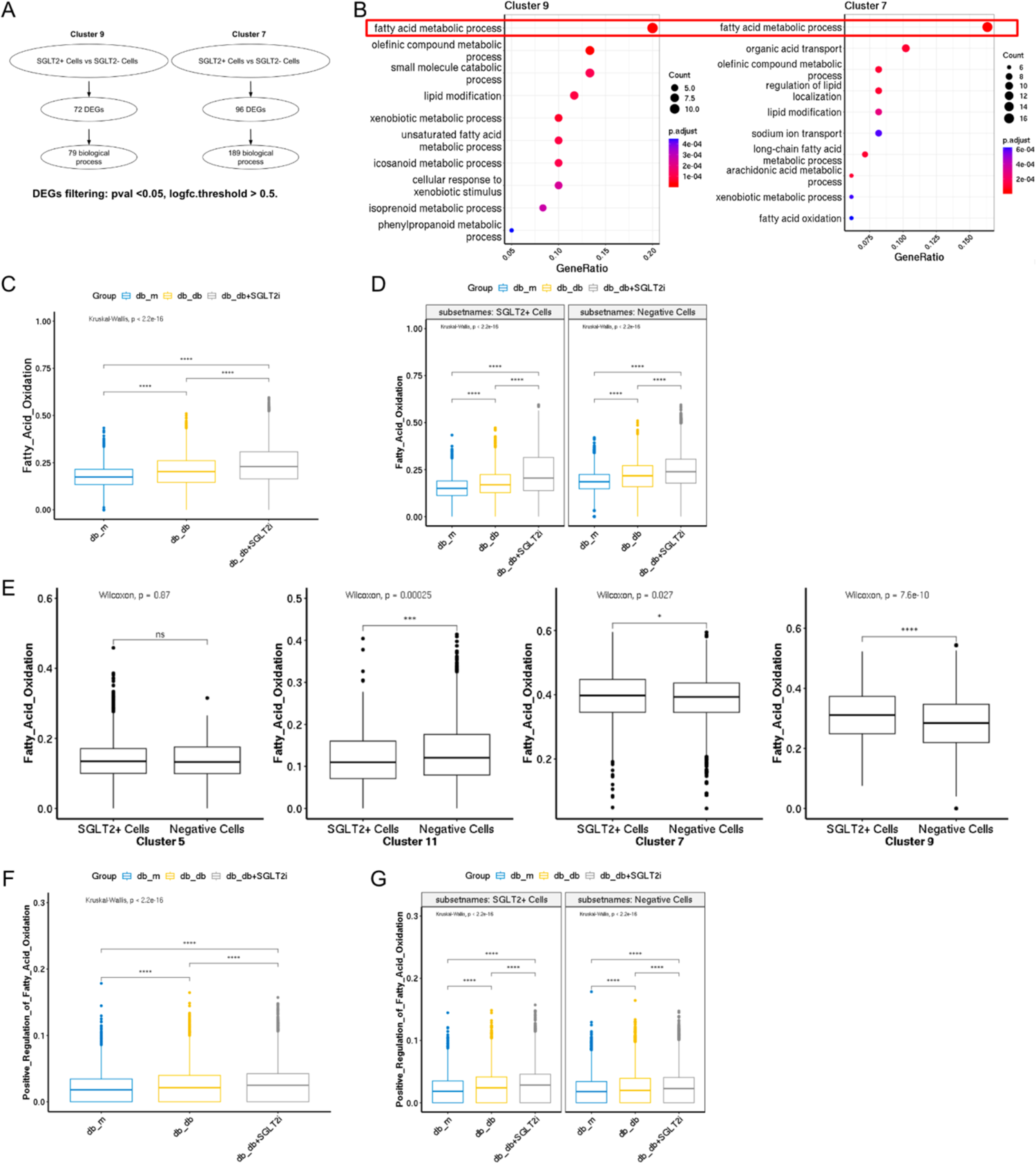
Fatty acid oxidation in diabetes. **(A),** Flowchart of mapping DEGs from clusters 9 and 7 for GO enrichment analysis. **(B),** Top 10 biological processes in clusters 9 and 7. **(C),** Signature scoring analysis of the fatty acid oxidation. **(D),** Signature scoring analysis of the fatty acid oxidation in SGLT2(+) vs. SGLT2(-) cells in groups. **(E),** Signature scoring analysis of the fatty acid oxidation in clusters 5, 11, 7, 9. **(F),** Signature scoring analysis of positive regulation of fatty acids oxidation in groups. **(G),** Signature scoring analysis of positive regulation of fatty acid oxidation in SGLT2(+) vs. SGLT2(-) cells in groups. All differences were analyzed by the Wilcox test. ns, P>=0.05. *, P < 0.05. ***, P < 0.001. ****, P < 0.0001.

Fatty acid oxidation includes three mechanisms: peroxisomal β-oxidation, mitochondrial β-oxidation, and microsomal μ-oxidation [29] (**Fig. 4A**). As shown in **Fig. 4B**, we computed the expression profile of fatty acid oxidation enzymes. Compared to the control group, diabetes increased expression of Cpt1a/2 and Abcd3, suggesting more fatty acid transporting to mitochondria and peroxisomes (**Fig.4A** and **4B**). Some enzymes of fatty acid oxidation were also elevated, including Acadvl, Hadha/b, Acaa2, Eci1, Ehhadh, Hsd17b4, Cyp4a10, Cyp4a12b, and the expression of Cyp4a14, while Acads, Acadm, Acadl, Eci2, Acox1 and Cyp4a12a were reduced (**Fig.4B**). These results indicate diabetes elevated fatty acid oxidation mostly through mitochondrial oxidation and peroxisomal oxidation. In the SGLT2i treatment group, most enzymes had higher expression compared to the diabetic group, except Cpt2, Acads, and Acadm, suggesting that SGLT2 inhibition increased the peroxisomal β-oxidation, microsomal χο-oxidation, and the long-chain fatty acid catalyzed in the mitochondria (**Fig.4A** and **4B**).

**Figure 4:**
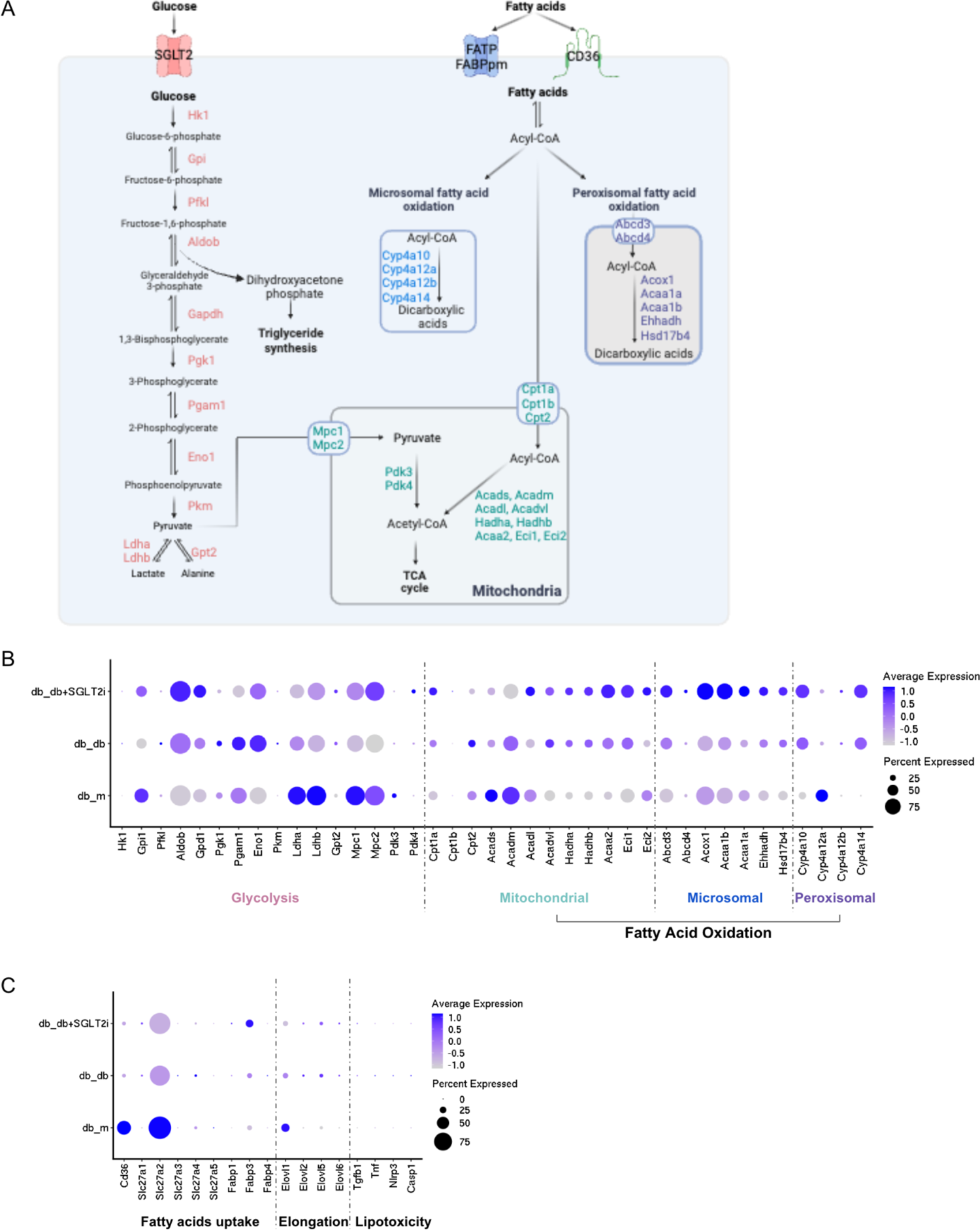
Regulation of fatty acid oxidation. **(A),** Pathways of glycolysis (32, 33) and fatty acid oxidation (34, 35). Created with BioRender.com. **(B),** Expression profile of genes regulating glycolysis and fatty acid oxidation. **(C),** Expression profile of genes regulating long-chain fatty acid uptake, elongation, and lipotoxicity. The dot color represents the expression level, and the legend is provided beside the plot. The dot size represents the % of cells in each cluster expressing the respective gene. SGLT2, sodium-glucose cotransporter 2; FATP, fatty acid transport protein; FABPpm, fatty acid binding protein; tricarboxylic acid (TCA) cycle.

Given our results suggesting the crucial mechanism of SGLT2 inhibition in long-chain fatty acid oxidation, we next investigate the source of long-chain fatty acids. By analyzing the transporters of long-chain fatty acids, we found that Cd36 and fatty acid transport protein 2 (Slc27a2) were expressed in over 50% of cells and had the highest expression in the control group (**Fig.4C**).

Diabetes or SGLT2i treatment did not increase the expression of these genes. However, the treatment group increased the expression of Fabp3, while diabetes slightly increased its expression compared to the control group. We further compared the expression of fatty acid elongases and showed that SGLT2i treatment did not elevate the elongation of fatty acid compared to the diabetic group while diabetes increased the expression of Elovl2, Elovl5, and Elovl6 in less than 10% of the cells and the control group had the highest expression of Elovl1 in 25% of the cells (**Fig.4C**). These results indicate that SGLT2i treatment did not increase the supply of long-chain fatty acids through cellular uptake or elongation. Similarly, SGLT2 inhibition did not aggregate the lipotoxicity induced by lipid accumulation via TNFα, TGFβ, and NLRP3 inflammasome pathways (29)(**Fig.4C**).

## Discussion

In the present study, we provide a comprehensive analysis of the kidney transcriptome in diabetic mice with SGLT2i treatment. We assessed the metabolic reprogramming in proximal tubular cells with or without SGLT2i treatment in SGLT2(+) vs. SGLT2(-) cells. Our results reveal that SGLT2i reduces gene expression consistent with a reduction in glycolysis and increases in the removal of glycolytic substrate accumulation in the mitochondria. SGLT2i treatment also upregulates genes related to fatty acid oxidation in diabetes on both SGLT2(+) vs. SGLT2(-) cells without inducing lipotoxicity.

It is known that SGLT2 inhibition protects against kidney injury in diabetes (30). As a major glucose transporter, glycolysis is important in studying the renoprotective mechanisms of SGLT2i. We found that SGLT2i alters gene expression consistent with diminished glucose influx with a higher rate of transport of pyruvate to mitochondria fueling the TCA cycle. These effects may contribute to restoring the glucose overload-induced substrate accumulation and metabolic dysfunction in tubular cells. Our findings are consistent with prior studies that showed reduced glycolysis in diabetes (23, 31). In addition, a mouse study elucidated the impaired glycolysis in diabetic mice by labeling with ^13^C-glucose characterized by increased glucose influx in diabetic mice (25). However, in this study, SGLT2i failed to reverse the dysfunction, which may be due to insufficient duration of treatment with SGLT2i.

Cai et al. reported a markedly higher lipid accumulation in the tubulointerstitial space of human diabetic kidney sections (31). This study also demonstrated that SGLT2 inhibition suppresses the metabolic transition from lipid oxidation to glycolysis using a mouse diabetic model and cultured tubular epithelial cells with upregulation of the palmitoyltransferase-1α (Cpt1a) and long-chain acyl-CoA dehydrogenase (Acadl). Our data demonstrate that SGLT2i primarily increases expression of genes consistent with long-chain fatty acids in mitochondrial oxidation, along with an overall higher level of microsomal fatty acid oxidation and peroxisomal fatty acid oxidation. Together with our data on glycolytic pathways and fatty acid lipotoxicity, SGLT2 inhibition may regulate the pathogenesis of diabetic kidney disease by reducing both lipotoxicity and glucotoxicity.

We observed a similar effect of SGLT2 inhibition on SGLT2(+) vs. SGLT2(-) cells, implicating a non-cell-autonomous effect. Given the limited numbers of SGLT2(+) cells in tubules, our data suggest that the SGLT2-independent effects of SGLT2 inhibition could occur via metabolic feedback or off-target effects to explain the consistency of significant changes in gene expression in both SGLT2(+) and SGLT2(-) cells observed between control and diabetic groups and between diabetic and diabetic + SGLT2i groups. It deserves further exploration in future studies. One possibility is that SGLT2i causes foundational changes in SGLT2(+) cells, and these are indirectly transmitted to SGLT2(-) cells. We cannot exclude that these main metabolic effects of SGLT2i occur independently of SGLT2 (e.g., through inhibition of another transporter, e.g., SGLT1). Distinguishing between these possibilities is an important future direction for this work.

The limitations of this work include the analysis of a single dose of SGLT2i, for a fixed duration, using a single class of SGLT2i. These were publicly available data from an established experiment with accompanying clinical and histologic data consistent with known benefits of SGLT2 inhibition. We also lacked metabolomic or histologic data from individual mice, but arguably, the ability to segregate effects in SGLT2(+) vs. SGLT2(-) cells is most readily available using single-cell transcriptomic data.

In summary, these results suggest that the reversal of diabetes-induced metabolic derangements in both SGLT2(+) and SGLT2(-) cells is crucial to the renoprotective effect of SGLT2i to reduce tubular injury. Inhibition of SGLT2 recovers metabolic dysfunction and renal injury induced by glucotoxicity and lipotoxicity in a non-cell autonomous fashion. Our study provides new insights into the beneficial effects of SGLT2 inhibition in diabetic kidney disease.

## Acknowledgments

Y.S. was supported by the Larry L. Hillblom Foundation. V.B. was funded, in part, by the NIH/NIDDK (R01 DK115770). The computing for this project was performed on the Sherlock cluster. We would like to thank Stanford University and the Stanford Research Computing Center for providing computational resources and support that contributed to these research results. We would like to thank the He Laboratory (Icahn School of Medicine at Mount Sinai) for sharing the scRNA-seq data set (26).

## Author contributions

Y.S. designed the study, performed all computational analyses, created the figures, and wrote the manuscript. V.B. supervised the project and revised the manuscript.

## References

1. Prevention CfDCa. National Diabetes Statistics Report. Centers for Disease Control and Prevention: US Dept of Health and Human Services; 2022.

2. Koyama AK, Cheng YJ, Brinks R, Xie H, Gregg EW, Hoyer A, et al. Trends in lifetime risk and years of potential life lost from diabetes in the United States, 1997-2018. PLoS One. 2022;17(5):e0268805.doi:10.1371/journal.pone.0268805

3. Greco DS, Broussard JD, Peterson ME. Insulin therapy. Vet Clin North Am Small Anim Pract. 1995;25(3):677–89.doi:10.1016/s0195-5616(95)50062-2

4. Hostalek U, Gwilt M, Hildemann S. Therapeutic Use of Metformin in Prediabetes and Diabetes Prevention. Drugs. 2015;75(10):1071–94.doi:10.1007/s40265-015-0416-8

5. Del Prato S, Pulizzi N. The place of sulfonylureas in the therapy for type 2 diabetes mellitus. Metabolism. 2006;55(5 Suppl 1):S20–7.doi:10.1016/j.metabol.2006.02.003

6. Rieg T, Vallon V. Development of SGLT1 and SGLT2 inhibitors. Diabetologia. 2018;61(10):2079–86.doi:10.1007/s00125-018-4654-7

7. Mosenzon O, Wiviott SD, Cahn A, Rozenberg A, Yanuv I, Goodrich EL, et al. Effects of dapagliflozin on development and progression of kidney disease in patients with type 2 diabetes: an analysis from the DECLARE-TIMI 58 randomised trial. Lancet Diabetes Endocrinol. 2019;7(8):606–17.doi:10.1016/S2213-8587(19)30180-9

8. Perkovic V, Jardine MJ, Neal B, Bompoint S, Heerspink HJL, Charytan DM, et al. Canagliflozin and Renal Outcomes in Type 2 Diabetes and Nephropathy. N Engl J Med. 2019;380(24):2295–306.doi:10.1056/NEJMoa1811744

9. Radholm K, Figtree G, Perkovic V, Solomon SD, Mahaffey KW, de Zeeuw D, et al. Canagliflozin and Heart Failure in Type 2 Diabetes Mellitus: Results From the CANVAS Program. Circulation. 2018;138(5):458–68.doi:10.1161/CIRCULATIONAHA.118.034222

10. Kato ET, Silverman MG, Mosenzon O, Zelniker TA, Cahn A, Furtado RHM, et al. Effect of Dapagliflozin on Heart Failure and Mortality in Type 2 Diabetes Mellitus. Circulation. 2019;139(22):2528–36.doi:10.1161/CIRCULATIONAHA.119.040130

11. Mather A, Pollock C. Glucose handling by the kidney. Kidney Int Suppl. 2011(120):S1–6.doi:10.1038/ki.2010.509

12. Bakris GL, Fonseca VA, Sharma K, Wright EM. Renal sodium-glucose transport: role in diabetes mellitus and potential clinical implications. Kidney Int. 2009;75(12):1272–7.doi:10.1038/ki.2009.87

13. Skrtic M, Yang GK, Perkins BA, Soleymanlou N, Lytvyn Y, von Eynatten M, et al. Characterisation of glomerular haemodynamic responses to SGLT2 inhibition in patients with type 1 diabetes and renal hyperfiltration. Diabetologia. 2014;57(12):2599–602.doi:10.1007/s00125-014-3396-4

14. Koh ES, Kim GH, Chung S. Intrarenal Mechanisms of Sodium-Glucose Cotransporter-2 Inhibitors on Tubuloglomerular Feedback and Natriuresis. Endocrinol Metab (Seoul). 2023;38(4):359–72.doi:10.3803/EnM.2023.1764

15. Chen X, Delic D, Cao Y, Shen L, Shao Q, Zhang Z, et al. Renoprotective effects of empagliflozin are linked to activation of the tubuloglomerular feedback mechanism and blunting of the complement system. Am J Physiol Cell Physiol. 2023;324(4):C951–C62.doi:10.1152/ajpcell.00528.2022

16. Ke Q, Shi C, Lv Y, Wang L, Luo J, Jiang L, et al. SGLT2 inhibitor counteracts NLRP3 inflammasome via tubular metabolite itaconate in fibrosis kidney. FASEB J. 2022;36(1):e22078.doi:10.1096/fj.202100909RR

17. Xu J, Kitada M, Ogura Y, Liu H, Koya D. Dapagliflozin Restores Impaired Autophagy and Suppresses Inflammation in High Glucose-Treated HK-2 Cells. Cells. 2021;10(6).doi:10.3390/cells10061457

18. Pirklbauer M, Sallaberger S, Staudinger P, Corazza U, Leierer J, Mayer G, Schramek H. Empagliflozin Inhibits IL-1beta-Mediated Inflammatory Response in Human Proximal Tubular Cells. Int J Mol Sci. 2021;22(10).doi:10.3390/ijms22105089

19. Feng L, Chen Y, Li N, Yang X, Zhou L, Li H, et al. Dapagliflozin delays renal fibrosis in diabetic kidney disease by inhibiting YAP/TAZ activation. Life Sci. 2023;322:121671.doi:10.1016/j.lfs.2023.121671

20. Gordin D, Shah H, Shinjo T, St-Louis R, Qi W, Park K, et al. Characterization of Glycolytic Enzymes and Pyruvate Kinase M2 in Type 1 and 2 Diabetic Nephropathy. Diabetes Care. 2019;42(7):1263–73.doi:10.2337/dc18-2585

21. Qi W, Keenan HA, Li Q, Ishikado A, Kannt A, Sadowski T, et al. Pyruvate kinase M2 activation may protect against the progression of diabetic glomerular pathology and mitochondrial dysfunction. Nat Med. 2017;23(6):753–62.doi:10.1038/nm.4328

22. Beitner R, Kalant N. Stimulation of glycolysis by insulin. J Biol Chem. 1971;246(2):500–3

23. Schaub JA, AlAkwaa FM, McCown PJ, Naik AS, Nair V, Eddy S, et al. SGLT2 inhibitors mitigate kidney tubular metabolic and mTORC1 perturbations in youth-onset type 2 diabetes. J Clin Invest. 2023;133(5).doi:10.1172/JCI164486

24. Li J, Liu H, Takagi S, Nitta K, Kitada M, Srivastava SP, et al. Renal protective effects of empagliflozin via inhibition of EMT and aberrant glycolysis in proximal tubules. JCI Insight. 2020;5(6).doi:10.1172/jci.insight.129034

25. Kogot-Levin A, Riahi Y, Abramovich I, Mosenzon O, Agranovich B, Kadosh L, et al. Mapping the metabolic reprogramming induced by sodium-glucose cotransporter 2 inhibition. JCI Insight. 2023;8(7).doi:10.1172/jci.insight.164296

26. Wu J, Sun Z, Yang S, Fu J, Fan Y, Wang N, et al. Kidney single-cell transcriptome profile reveals distinct response of proximal tubule cells to SGLT2i and ARB treatment in diabetic mice. Mol Ther. 2022;30(4):1741–53.doi:10.1016/j.ymthe.2021.10.013

27. Andreatta M, Carmona SJ. UCell: Robust and scalable single-cell gene signature scoring. Comput Struct Biotechnol J. 2021;19:3796–8.doi:10.1016/j.csbj.2021.06.043

28. Ross BD, Espinal J, Silva P. Glucose metabolism in renal tubular function. Kidney Int. 1986;29(1):54–67.doi:10.1038/ki.1986.8

29. Yuan Q, Tang B, Zhang C. Signaling pathways of chronic kidney diseases, implications for therapeutics. Signal Transduct Target Ther. 2022;7(1):182.doi:10.1038/s41392-022-01036-5

30. Vallon V, Verma S. Effects of SGLT2 Inhibitors on Kidney and Cardiovascular Function. Annu Rev Physiol. 2021;83:503–28.doi:10.1146/annurev-physiol-031620-095920

31. Cai T, Ke Q, Fang Y, Wen P, Chen H, Yuan Q, et al. Sodium-glucose cotransporter 2 inhibition suppresses HIF-1alpha-mediated metabolic switch from lipid oxidation to glycolysis in kidney tubule cells of diabetic mice. Cell Death Dis. 2020;11(5):390.doi:10.1038/s41419-020-2544-7

32. Hay N. Reprogramming glucose metabolism in cancer: can it be exploited for cancer therapy? Nat Rev Cancer. 2016;16(10):635–49.doi:10.1038/nrc.2016.77

33. Zangari J, Petrelli F, Maillot B, Martinou JC. The Multifaceted Pyruvate Metabolism: Role of the Mitochondrial Pyruvate Carrier. Biomolecules. 2020;10(7).doi:10.3390/biom10071068

34. He Q, Chen Y, Wang Z, He H, Yu P. Cellular Uptake, Metabolism and Sensing of Long-Chain Fatty Acids. Front Biosci (Landmark Ed). 2023;28(1):10.doi:10.31083/j.fbl2801010

35. Kersten S. Integrated physiology and systems biology of PPARalpha. Mol Metab. 2014;3(4):354–71.doi:10.1016/j.molmet.2014.02.002

